# Spatial metrics of tumour vascular organisation predict radiation efficacy in a computational model

**DOI:** 10.1101/029595

**Authors:** Jacob G. Scott, Alexander G. Fletcher, Alexander R.A. Anderson, Philip K. Maini

## Abstract

Intratumoural heterogeneity is known to contribute to poor therapeutic response. Variations in oxygen tension in particular have been correlated with changes in radiation response *in vitro* and at the clinical scale with overall survival. Heterogeneity at the microscopic scale in tumour blood vessel architecture has been described, and is one source of the underlying variations in oxygen tension. We endeavour to determine whether histologic scale measures of the erratic distribution of blood vessels within a tumour can be used to predict differing radiation response. Using a two-dimensional hybrid cellular automaton model of tumour growth, we evaluate the effect of vessel distribution on cell survival outcomes of simulated radiation therapy. Using the standard equations for the oxygen enhancement ratio for cell survival probability under differing oxygen tensions, we calculate average radiation effect over a range of different vessel densities and organisations. We go on to quantify the vessel distribution heterogeneity and measure spatial organization using Ripley’s *L* function, a measure designed to detect deviations from spatial homogeneity. We find that under differing regimes of vessel density the correlation coefficient between the measure of spatial organization and radiation effect changes sign. This provides not only a useful way to understand the differences seen in radiation effect for tissues based on vessel architecture, but also an alternate explanation for the vessel normalization hypothesis.

**Author Summary:** In this paper we use a mathematical model, called a hybrid cellular automaton, to study the effect of different vessel distributions on radiation therapy outcomes at the cellular level. We show that the correlation between radiation outcome and spatial organization of vessels changes signs between relatively low and high vessel density. Specifically, that for relatively low vessel density, radiation efficacy is decreased when vessels are more homogeneously distributed, and the opposite is true, that radiation efficacy is improved, when vessel organisation is normalised in high densities. This result suggests an alteration to the vessel normalization hypothesis which states that normalisation of vascular beds should improve radio- and chemo-therapeutic response, but has failed to be validated in clinical studies. In this alteration, we provide a metric that differentiates between vascular architectures in different density regimes in which the hypothesis holds and does not and that could be used for quantitative histologic analysis of tumours, and for radiation dose personalisation.

**Author Contributions:** Conceived and designed the experiments: JGS, AGF, ARAA, PKM

Performed the experiments: JGS

Analyzed the data: JGS, AGF, ARAA, PKM

Wrote the paper: JGS, AGF, ARAA, PKM

## Introduction

It is increasingly recognised that an important aspect of cancers is their heterogeneity [1]. This heterogeneity exists between patients, between different tumours within a single patient [2], within the differing cellular populations in a single tumour and even at the genetic scale between cancer cells originating from the same ancestor [3]. In particular, microenvironmental heterogeneity is becoming widely accepted as a key factor in tumour progression and response to therapy [1]. Nutrients, growth factors, extracellular matrix and other cell types are all part of the normal tissue that surrounds and pervades a solid tumour and has been shown to vary widely across different tumour stages and types. This is, in part, due to the dynamic and heterogeneous interplay between the tumour and its microenvironment.

Radiation biologists have, for many years, understood the importance of cell biological and microenvironmental factors on radiation response. Current radiation therapy dose planning, however, largely neglects this information and is, instead, based on years of clinical experience using intuition and trial and error. As such, there remains limited tailoring of dose planning to an individual patient’s tumour. With the advent of modern quantitative histologic [4] and biological imaging methods [5], however, this paradigm is poised to change.

Research in this area over the last decade [6] has sought to understand the macroscopic spatial distribution of hypoxia within tumours using non-invasive imaging. This information has then been utilised to develop spatially heterogeneous dose plans to improve tumour control. For example, Malinen et al. [7] inferred average oxygen concentrations from radiocontrast concentrations measured by Dynamic Contrast Enhanced (DCE) Magnetic Resonance Imaging (MRI) in a dog sarcoma. Other work to understand the effects of radiation in individual patients has utilized MRI scans in combination with mathematical models of tumour growth. These models have incorporated heterogeneity in cell type by considering a two compartment spatial partial differential equation (PDE) model, separately considering proliferation and motility, without consideration of oxygen effects [8] to explain different radiation responses measured by changes in tumour size over time. More recently, cellular automaton models of stem cell driven tumours, comprised of populations with differences in proliferative phenotype [9] were used to compare tumour response to a range of spatially heterogeneous radiation dose plans or to different orders of radio-and chemo-therapeutic strategies [10]. What is lacking to date, however, is research into how tissue level microenvironmental heterogeneity can affect radiation response, and how this could be inferred from patient data.

Heterogeneity in tumour oxygenation, in particular the occurrence of hypoxia, is a well-known cause of radiation therapy failure [11, 12]. Work done *in vitro* to understand differences in radiation effect due to oxygenation differences have been valuable, and have established an empiric relationship called the Oxygen Enhancement Ratio (OER) [13] to understand how radiation efficacy varies with oxygen concentration. These studies do not, however, allow us to understand the effects of radiation *in vivo,* as they do not consider the heterogeneity of oxygenation at the microscopic, cellular scale.

Beyond the macroscopic changes in oxygenation, vessel and cellular density and metabolism, a number of theoretical studies have suggested that the local microscopic heterogeneity in oxygenation can vary widely in space and time in tumours [14, 15, 16] and healthy tissues [17] alike. In addition a large body of work has sought to understand an apparent paradox of therapy directed at angiogenesis, the process of new vessel creation [18]. In short, it was thought that by blocking a cancer’s ability to create new vessels, it would be possible to starve the cancer of nutrients and oxygen, quickly leading to its demise. While effective anti-angiogenesis drugs have been developed, this promise never came to fruition. The leading hypothesis to explain this failure is termed the ‘vascular normalization hypothesis’ [19], which suggests that these drugs, which block vascular endothelial growth factor (VEGF), do not simply inhibit new vessel production, but instead work by normalizing the vascular bed in question by pruning out ineffective vessels. This hypothesis, while well supported by experimental work, has not yet been able to fully explain results at the clinical scale [20]. These studies highlight an opportunity to improve our understanding of how spatio-temporal oxygen dynamics at the cellular scale affect tissue-level response to therapy. To this end, we develop a computational, hybrid cellular automaton model of a tumour growing within a surrounding normal tissue in a vascularized domain which we use to investigate whether spatial statistics gleaned from measures of vascular organisation can be used to predict radiation efficacy. By identifying broad relationships between *in silico* tissue architecture and radiation response, we aim to progress toward a translatable method of radiation plan optimization using information extracted from biopsies.

The remainder of this paper is structured as follows. We first elucidate our methods by describing the underlying rules and parameters governing the cellular automaton as well as the method of calculating oxygen transport and uptake. In the results section we describe simulation results concerning healthy tissue growth in regularly arranged and then heterogeneous vascular architectures. We then describe tumour growth and invasion in regular architectures and follow this with observations concerning the distribution of oxygen tension in different possible vascular organisations. We then develop a metric, based on Ripley’s *L* function [21], which allows us to quantify and correlate these patterns with radiation response. We end with a discussion of how this metric reconciles some of the difficulties with the vascular normalisation hypothesis, and suggest how the metric might be used in the clinic to personalise radiation dose planning.

## Methods

We consider the effect of a heterogeneous microenvironment through the inclusion of *en face* blood vessels modeled as point sources of oxygen, and through competition of tumour cells with an initial field of healthy cells. We begin by creating a hybrid cellular automaton (HCA) model [22] in which we describe cells by individual agents whose states are updated over synchronous discrete time steps of fixed duration on the timescale of the cell cycle (automaton time steps), and which occupy sites on a two-dimensional square lattice representing a slice of tissue. The size of the lattice spacing is chosen such that each automaton element is approximately the size of a single cell. A second, identical lattice is created on which we approximate the continuous concentration of a freely diffusible molecule, representing oxygen which is updated on a finer timescale (oxygen timestep - see supplemental information for further description of the relationship between these timescales). While the cells and oxygen distribution are updated on separate lattices, each influences the other. The feedback between the cells and the microenvironment is captured through a partial differential equation (PDE) governing oxygen transport and consumption. Although the parameter values in our model are drawn from data on a particular cancer type, the primary brain tumour glioblastoma, most of the underlying model assumptions are likely to apply to many other vascularized solid tumours.

### Cellular automaton model of cell behaviour

Cell fate decisions in our model are determined by a number of microenvironmental and cell state-specific thresholds and values. The order in which cells are chosen to decide their fate is computed in a random fashion so as to avoid any order bias. To determine the position of cells within our domain, we tessellate the continuous domain on which the PDE is defined into squares of size Δ*x* × Δ*x* to arrive at a regular lattice occupied by both cells and vessels. While all cells are assumed to be the same size and shape (a single lattice site), we model two cell types, cancerous and healthy cells, each of which can divide to produce two identical daughters. The probabilities and thresholds for these cell fate decisions are listed in Table 1 and Fig. 1, respectively, and will be discussed individually in the coming sections.

**Table 1.**
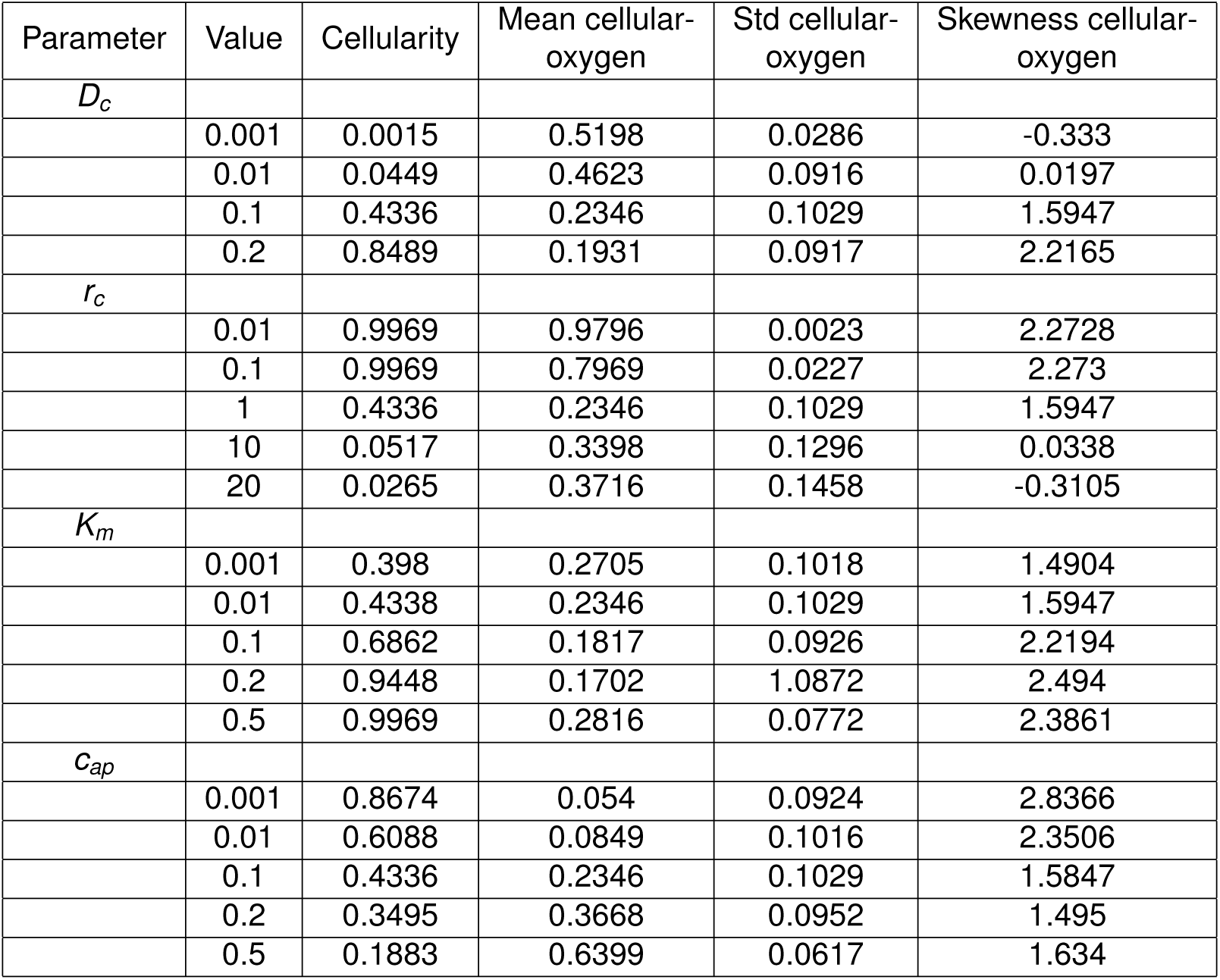
Dimensional model parameters and their estimates.

**Figure 1.**
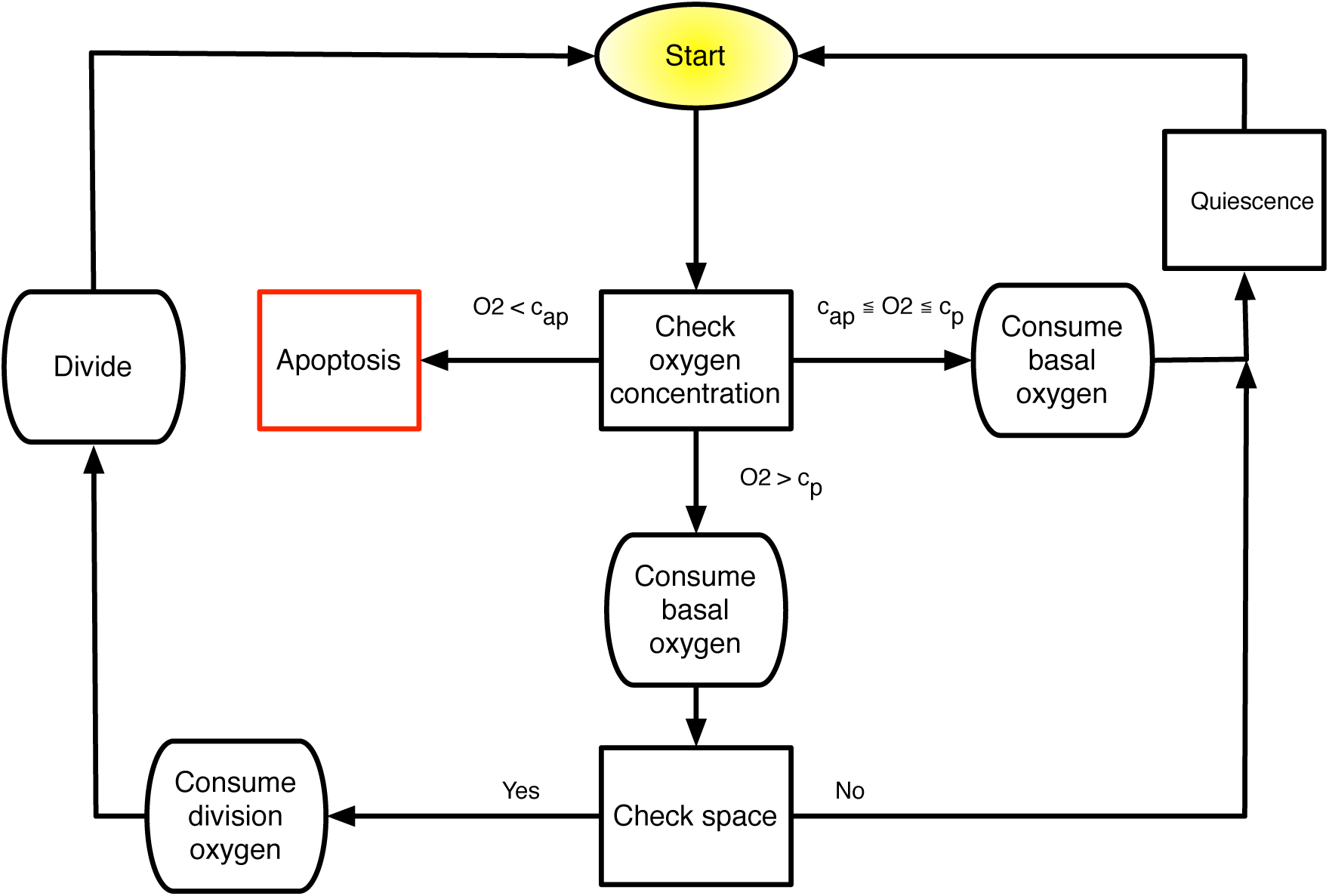
Schematic of discrete time updating algorithm for the HCA. At each cellular automaton update, each cell in the domain undergoes a series of fate decisions based on intrinsic cell parameters and microenvironmental cues.

**Oxygen consumption and transport.** With the advent of single-cell techniques for measuring oxygen consumption rates, quantitative data concerning the differential oxygen consumption rates of different types of cancer cells are now available. It is widely accepted that, in general, cancer cells are more metabolically active than healthy cells because of the Warburg shift [23] or other metabolic abnormalities, and we thus use a modification of normal consumption.

The continuous component of our model describes the distribution and consumption of nutrients. While it is clear that many nutrients are of biological importance, in this model we focus on oxygen as it is central to the DNA-damaging effects of photon radiation therapy [24]. Blood vessels, which are each modeled as point sources of oxygen, occupying one lattice site, are placed randomly throughout the lattice at the start of a given simulation, at a specified spatial density ⊝. While in reality, vessels can be different sizes, they tend to be normally distributed around a mean size, and a variety of metrics including number of vessels, total vessel area and vascular fraction have been used interchangeably [25, 26]. As we are interested in the dynamics over the time scale of radiation treatment (order days), in all simulations we neglect vascular remodeling and angiogenesis, processes which are triggered by hypoxia and mediated by hypoxia inducible factor-1*α* (HIF-1*α*) and VEGF, among other molecules, and have been well studied [27, 28]. Each vessel is assumed to carry an amount of oxygen equal to that carried in the arterial blood [29]. This oxygen is then assumed to diffuse into the surrounding tissue and be consumed by normal and tumour cells.

The spatiotemporal evolution of the oxygen field is described by the reaction-diffusion PDE

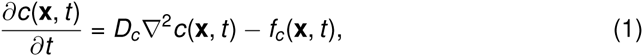
where *c*(**x**, t) is the concentration of oxygen at a given time *t ≥* 0 and position **x** ∈ (0, *NΔx*)^2^\ V, where Δ*x* is the length of a typical cell, *N* is the number of lattice sites on a side of the *N × N* domain and V = {**x**_1_, **x**_2_,…, **x***_v_*} denotes the set of points occupied by blood vessels, which are *v* in number. We define *D_c_* to be the diffusion coefficient of oxygen, which for simplicity we assume to be constant, corresponding to linear, isotropic diffusion. The function *f_c_*(**x**, *t*) denotes the cellular uptake of oxygen, whose form is given by

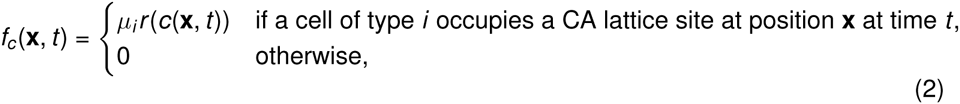
where *i ∈* {*H, T*} signifying Health (*H*) and Tumour (*T*) cells, respectively. Here *μ_i_* is defined as the cell type-specific oxygen consumption rate constant which modulates *r*(*c*), the oxygen-dependent consumption rate, which we assume to have Michaelis-Menten form

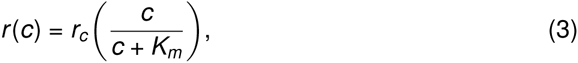
where *r_c_* and *K_m_* denote the maximal uptake rate and effective Michaelis-Menten constant, respectively. We supplement equation (1) with the following initial and boundary conditions. We begin with a spatially uniform oxygen distribution *c*(**x**, 0) = *c*_0_, a positive constant, with the entire domain, (0, *N*Δ*x*)^2^\ V, occupied by normal cells. To simulate carcinogenesis, we replace the cell in the central lattice site with a tumour cell. Vessels are placed throughout the domain at a prescribed density and pattern, depending on the situation being simulated. To approximate the constant value of the oxygen concentration in the arterial blood we impose *c*(**x**, *t*) = *c*_max_ at each point, ***X*** ∈ ***V***, where there is a vessel. We further impose no-flux boundary conditions at the edge of the domain,

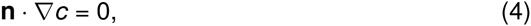
where **n** is the unit normal outward vector to the domain boundary.

**Proliferation.** Cell proliferation is governed by a complicated set of subcellular processes, and has been modelled in great detail [30, 31]. The cell cycle has many levels of complexity including checkpoints, specific temporal sequences and many biophysical processes that alter the cell shape and response to microenvironmental cues and stresses at different times in the cycle [32]. As we are interested in a phenomenon occurring at the much larger, tissue, scale, we will encapsulate this complexity into a simple rule set and threshold values on microenvironmental (oxygen) conditions and neighbourhood occupation.

For cells to progress through the cell cycle and divide, we require their local oxygen concentration to exceed a fixed threshold value *c*_p_, and that the neighbourhood (in this case utilizing a Moore neighbourhood) contains at least one empty space. If these constraints are met, then we allow the cell to divide with type-specific probability *pj* (where *j ∈* {*H, T*}) at each automaton time step, and place the resultant daughter in a randomly chosen empty neighbouring space. If no empty neighbouring space exists, then the cell becomes quiescent (see below). In the case of cancer cells, we relax this requirement to allow proliferation into healthy tissue, that is that normal cells are considered empty spaces when assaying for empty space for division.

**Cell death.** The main mechanisms of cancer cell death considered in this study occur due to severe hypoxia and radiation therapy. Healthy cells can also die through if they are selected for replacement by dividing cancer cells, a situation which would normally be ascribed to a cancer cell’s greater fitness, allowing it to out-compete healthy cells for space or nutrients locally, or possibly secondary to an acid-mediated event [23, 33].

Cells under extreme hypoxic conditions are often found to undergo autophagy (directly translated as ‘self-eating’), a state in which they become resistant to nutrient starvation [34], and cells are known to die over different time scales and by different mechanisms (apoptosis versus necrosis) depending on the magnitude and duration of the hypoxic insult. While these differences have been shown to affect tumour growth [35], as this is not the main focus of this model, we will simplify this scenario by assigning a rate, *p*_d_, for cell death at each cellular automaton update, when under extreme hypoxia (i.e. *c < c*_ap_ where *c*_ap_ ∈ (0, *c*_max_] is a fixed, model-specific positive constant).

To model the effect of radiation on heterogeneously oxygenated tissues, we begin with the linear-quadratic model of cell survival:

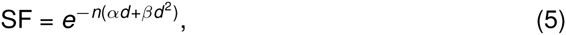
where *α* and *β* are radiobiologic parameters associated phenomenologically with cell kill resulting from ‘single hit’ events (*α*) and ‘double hit’ events (*β*) respectively, and *n* represents the number of doses of *d* Gy of radiation. This metric, while originally derived to fit *in vitro* data [36], has since been widely used to also describe the clinical efficacy of radiation [37, 38].

The basic interaction driving DNA damage from radiation is mediated either by free electrons or by free radicals formed by the interaction of photons with water. The DNA damage instantiated by these mechanisms can be repaired by reduction by locally available biomolecules containing sulfhydryl groups [24]. This damage can be made more permanent, however, by the presence of molecular oxygen, which binds with the DNA radical, forming a ‘non-restorable’ organic peroxide. This damage ‘fixation’ by oxygen creates a situation in which the damage now requires enzymatic repair, which occurs on a much longer timescale [24]. Regardless of the mechanism of radiation sensitisation by oxygen, the effect is well documented, and has been modelled by replacing the parameters *α* and *β* in equation (5) with functions of the oxygen concentration using the oxygen enhancement ratio (OER):

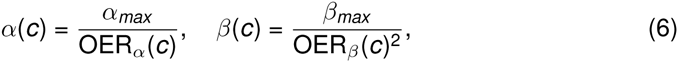
where *α_max_* and *β_max_* are the values of *α* and *β* under fully physiologically oxygenated conditions and *α*(c), *β*(c) are, respectively, the values of *α*, *β* at oxygen concentration *c*. We define the OER as a function of the oxygen concentration by using the empirically established relation [13, 39]

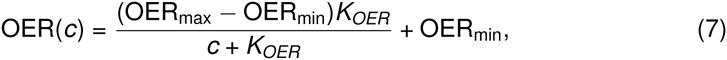
which is solved individually for *α* and *β* (see Table 1 for parameters).

**Quiescence.** In response to a lack of oxygen or other nutrients or external mechanical cues indicating overcrowding, cells enter a temporary state of quiescence during which no division occurs [40]. We model this process through an oxygen threshold (*c* < *c*_p_ where *c*_p_ ∈ [*c*_ap_, *c*_max_] is a fixed, model-specific positive constant) below which cellular division is not possible and by the spatial constraint that the cell is quiescent if there are no empty neighbouring lattice sites (using a Moore neighbourhood). In the case of a cancer cell, we require that at least one space be empty or inhabited by a normal cell, a situation in which we assume that the cancer cell will replace the normal cell if chosen for division. When the conditions for quiescense are no longer present, the cell’s state of quiescence is reversed.

**Normal tissue approximation.** The invasion of cancer cells into healthy tissue has been modelled extensively [41, 42, 43, 44]. While this process is not the focus of our study, we nevertheless must consider the healthy tissue surrounding our growing tumour, as it plays an important role in modulating the local oxygen concentration. Previous investigations have modelled tumour spheroid growth [45], or assumed a constant influx of oxygen from the boundary of the tissue, but as we endeavour to understand how heterogeneous vascularisation affects the growth of a tumour, we must also incorporate normal tissue effects [14]. To do this, we assume that the entire tissue is initially comprised of normal cells, which can divide if they have sufficient space, which consume oxygen at a rate of *μ_H_r*(*c*) and are subject to hypoxic cell death. In addition, normal cells can die through competition with cancer cells.

**Cell competition.** The ability to invade into normal tissue is one of the ‘Hallmarks of Cancer’ [46]. Several mechanisms for this have been proposed and modelled, including degradation of extracellular matrix by secreted proteases [41, 47] and mediated by excreted lactic acid [42, 43]. As this specific interaction is not our focus, we make the simplifying assumption that normal cells do not spatially inhibit cancer cell proliferation, and a cancer cell can freely proliferate and place a daughter onto a lattice location where a normal cell is located, causing the normal cell to die, either through an acid-mediated or simple fitness based competition mechanism [48]. Normal cells, however, are spatially inhibited by cancer cells and other normal cells.

**Vascular dynamics.** While in healthy tissue vessels are normally patent, as tumours grow the vasculature can experience significant spatio-temporal changes. There continues to be significant theoretical work to describe angiogenesis, the process of new vessel formation [22, 27], as well as the biophysical processes governing vascular occlusion [49, 50]. In this study, we seek only to understand the effect of heterogeneous oxygen concentrations on tumour growth and progression [14]. Therefore, we consider only the simplest scenario, in which there is no feedback from the environment to the vessels.

### Coupling models in the hybrid cellular automaton

Having described each of the constituent parts of the model, we now describe how they are coupled to form the HCA. We consider a two-dimensional lattice of size *N × N*, with each lattice element identified by coordinates (*i*Δ*x*, *j*Δ*x*) where *i*, *j ∈* {1,2,…, *N*}.

As schematised in Fig. 1, at each automaton time step every cell (chosen in random order) in the domain is subject to a series of decisions based on the current automaton state and local oxygen concentration, *c*(**x**, *t*). The specific checks that each cell undergoes, regardless of its phenotype, are:

1. compare *c*(**x**, *t*) to *c*_ap_ and *c*_p_;
2. consume oxygen at rate *μ_i_r_c_*, die, or remain quiescent;
3. check number of free neighbouring lattice sites;
4. determine proliferative behaviour: if proliferative constraints are met (space and oxygen), then consume additional oxygen for proliferation [51] and place daughter cell in randomly chosen empty neighboring lattice site, otherwise become quiescent.

**Cell-microenvironment interaction.** As we endeavour to understand the effects of a heterogeneous oxygen distribution on healthy tissues and tumours, we must consider a number of parameters that govern a cell’s behaviour in response to these environmental cues. Specifically, we consider the hypoxia threshold, *c*_ap_, below which cells will die, and be removed from the system. As cell fate decisions are made on a much slower time scale than oxygen diffusion, there is a potential for update bias which would entail cell fate decisions made long after the conditions for these decisions were met. To reduce this, we will ascribe a probability per automaton update, *p*_d_, to this death and removal, and update the HCA more frequently than the cell cycle time. We further consider a baseline oxygen consumption required for cellular processes, *r*_c_, and a threshold for cells to be proliferatively active, *c*_p_ [51].

**Non-dimensionalisation and parameter estimation.** In order to place the model in an appropriate spatial and temporal scale, we non-dimensionalise the system. We rescale time by a typical cell cycle time, τ = 16h [52], and lengths by a typical diameter Δ*x* = 50*μ*m for a glioma cell [53]. We choose to rescale the oxygen concentration to reflect the average oxygen concentration in the arterial blood, estimated at about 80mmHg (5.14 × 10^−13^ mol^−1^ cell^−1^) [54], so that the non-dimensional oxygen concentration at a vessel takes the value 1.

Introducing the non-dimensional variables 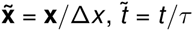 and 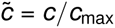, we define the new non-dimensional parameters

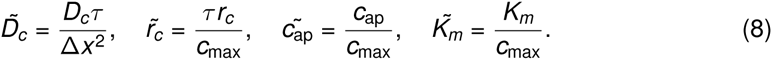

For notational convenience, we henceforth refer to the non-dimensional parameters only and drop the tildes for the aforementioned parameters. See Table 1 for a full list of dimensional parameter estimates and corresponding non-dimensional values.

**Numerical solution.** To solve equation (1) numerically, we discretize space and time by letting *t_k_* = *k*Δ*t*, where *k ∈* ℕ, and *x_i_,* = *i*Δ*x* and *yj* = *j*Δ*x* where *i*, *j ∈* {1,…, *N*} encode a square lattice of size *N × N*, and Δ*t* is the oxygen update time step. We approximate the oxygen concentration at time *t_k_* and position *x_i_*, *y_j_* by 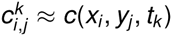. We use a central difference approximation for the Laplacian and thus approximate equation (1) by

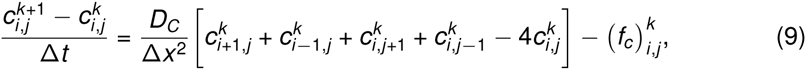
where 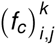 is the cell-specific oxygen consumption rate defined in equation (2). Rearranging equation (9) to obtain a solution for 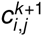 gives

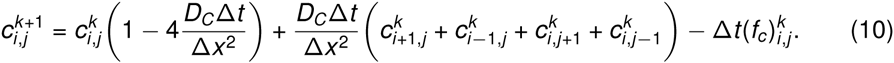

At each oxygen time step, the oxygen concentration in a given lattice site is updated using equation 10. To impose the zero-flux boundary conditions we modify equation (10) in the cases where *i*, *j* ∈ {1, *N*}. For example, to calculate the oxygen concentration experienced by a cell at the left-hand boundary (i.e. *i* =1) from outside the domain (i.e. *i* = 0) we discretize the no-flux boundary condition to obtain 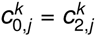, which we substitute into equation (10). Note that lattice sites occupied by vessels have fixed oxygen values (1 in the non-dimensional system), so we do not update them.

**Tumour control probability.** The tumour control probability (TCP) is a measure that approximates the probability of eradicating all cells within a tumour and thereby controlling (in this case ‘curing’) it. While a number of methods, both deterministic and stochastic, have been proposed to define such a measure [60], the most commonly used formulation assumes that the number of surviving cells capable of proliferation after radiation follows Poisson statistics and that the TCP is calculated by

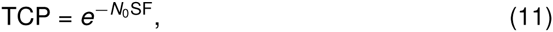
where *N*_0_ is the number of cells in the tissue before radiation therapy and *SF* is the surviving fraction defined by (5).

To compare the effect of radiation on different populations of cells in our model, we calculate the total number of cells surviving a given therapeutic intervention. To this end, we consider the individual survival probability of cells experiencing a given oxygen concentration. Let 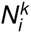 be the number of cells in the tissue that experience a oxygen concentration in the interval [*i*Δ*c,* (*i* + 1)Δ*c*) at time *t_k_* = *k*Δ*t*, for *i* ∈ {0,…, *M*}. Here we define Δ*c* = 0.01, and *M*Δ*c* = 1, reflecting the fact that the oxygen concentration is bounded by its non-dimensional value in the arterial blood, which is 1. We assume that radiation effect is instantaneous, and represent the time after radiation as *t_k_*_+1_. The number of cells experiencing an oxygen concentration in the interval [*i*Δ*c*, (*i* + 1)Δ*c*) immediately following a single radiation dose *d* is given by

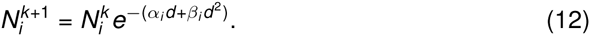

Thus the total number of surviving cells at time *t_k+_*_1_ is given by

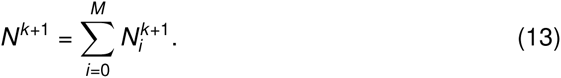

To calculate the number of surviving cells in a given domain after simulated radiation, we first calculate the individual values of *α_i_* and *β_i_* for each interval of oxygen concentration using equations (6) - (7). We can then compute the surviving fraction after a simulated single 2Gy dose of radiation (*d* = 2Gy, *n* = 1) by utilising equations (12) and (13) and assuming initial distribution of cellular-oxygen given by the simulations represented in Fig. 5.

## Results

### Oxygenation of healthy tissue

#### Regular vascular patterning preserves oxygen distribution shape but not mean

We first model a domain seeded with only healthy cells, and regular spatial distributions of vessels of varying density. Estimates in the literature for ‘normal’ vascular density span several orders of magnitude and are reported using a number of different measures [25, 61, 62], so we choose to calculate the actual physiological vascular density of this model by exploring regular vessel spacing and healthy tissue only. For each vessel spacing, the domain size is changed to accommodate the full complement of 40 vessels. In each of these cases, the pattern of the vascular distribution will remain constant, but the vascular density (number of vessels / domain size) will change. For ease of visualisation, we have chosen to plot the domains as equivalent sizes (see Fig. 2), and report the changing density.

**Figure 2.**
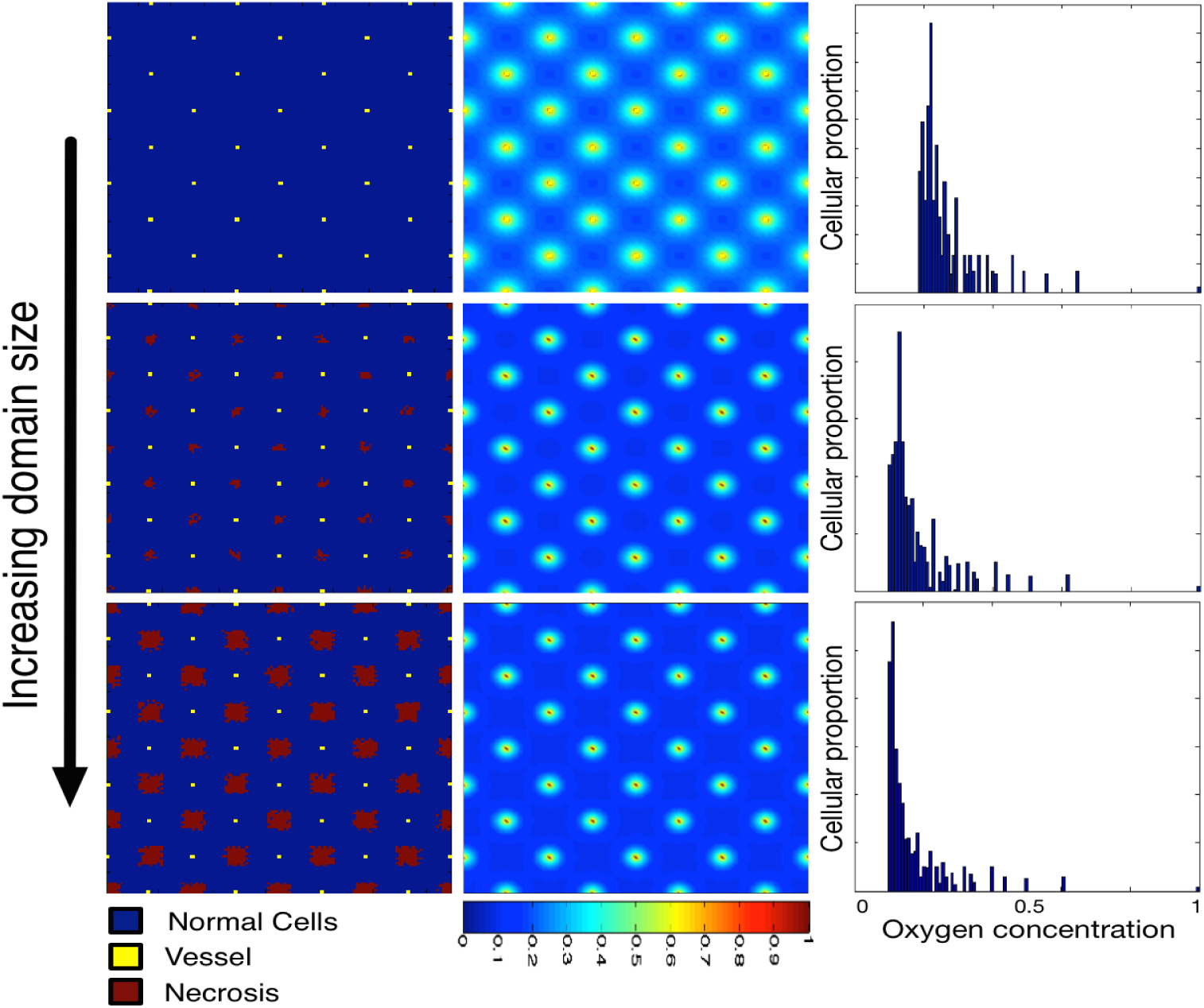
Varying the vascular density with regular spacing affects the carrying capacity and cellular-oxygen distribution in normal tissue. We plot healthy tissue growth and maintenance as we vary the vessel density (decreasing density from top to bottom ⊝ = 0.0031, 0.0027 and 0.0024). We plot cellular distributions (left) with associated spatial oxygen concentration (middle) and non-spatial distribution of cells versus oxygen concentration (right). These plots represent the system at dynamic equilibrium, in which cell death and birth is balanced across the tissue. The mean of the cellular-oxygen distribution decreases with vessel density (0.26, 0.18 and 0.16, top to bottom) while the standard deviation and skewness stay approximately constant (std = 0.09,0.1 and 0.1, skewness = 3.18,3.23 and 3.32). Domains are visualised as the same size for ease of comparison, but increase in size from top to bottom (domains are of size *N* × *N* where from top to bottom *N* = 114, 122, 130).

As expected, the closer the vascular packing in the domain, the more cells can be supported and the higher the mean oxygen tension (Fig. 2). There is a critical value for vessel density below which cells begin to die, and therefore the proportion of the domain inhabited by cells (termed carrying capacity henceforth) drops below unity. This value is between 2.7 × 10^−3^ vessels per lattice site and 3.1 × 10^−3^ vessels per lattice site for the parameters modeled (Table 1), which is well within an order of magnitude to the vascular area to tumour area ratio reported by Zhang et al. [63].

In addition to the carrying capacity, we investigate the distribution of cellular-oxygen concentration. This distribution provides a non-spatial summary statistic of an otherwise spatial entity: we can understand how many (or what proportion) of cells within the domain exist at certain oxygen concentrations, a measure which will become important later when we consider the effect of radiation. In Fig. 2 we note that, while the mean cellular oxygen concentration (mean of the cell-oxygen distribution) varies considerably (from a non-dimensional value of 0.16 to 0.45), the second and third moments of the distribution vary little, with the standard deviation staying between 0.08 – 0.11 and the skewness between 3.18 – 3.3. That is, the distribution of cellular oxygen concentration maintains its variance about its mean as well as its level of symmetry while the vascular patterning is regular.

#### Irregular vascular patterning results in variation in oxygen distribution shape

We next explore how the oxygen concentration that the population of healthy tissue supported experiences based on changes in density and patterning of vessels in irregularly vascularized domains. While this is a situation that would not likely be seen in the healthy state, understanding the behaviour of the model in this simple situation is valuable before progressing to more complex states. To this end we simulate the resulting healthy tissue growth and maintenance in a variety of randomly generated vascular patterns of the same density (Fig. 3). We plot the final, stable distribution of healthy cells for a random placement of vessels for a given density (⊝ = 0.001, top; ⊝ = 0.0025, middle; and ⊝ = 0.004, bottom) in the left and middle columns. In the right column, we plot the average distribution of cells at specific oxygen concentrations over ten simulations with different vascular patterns.

**Figure 3.**
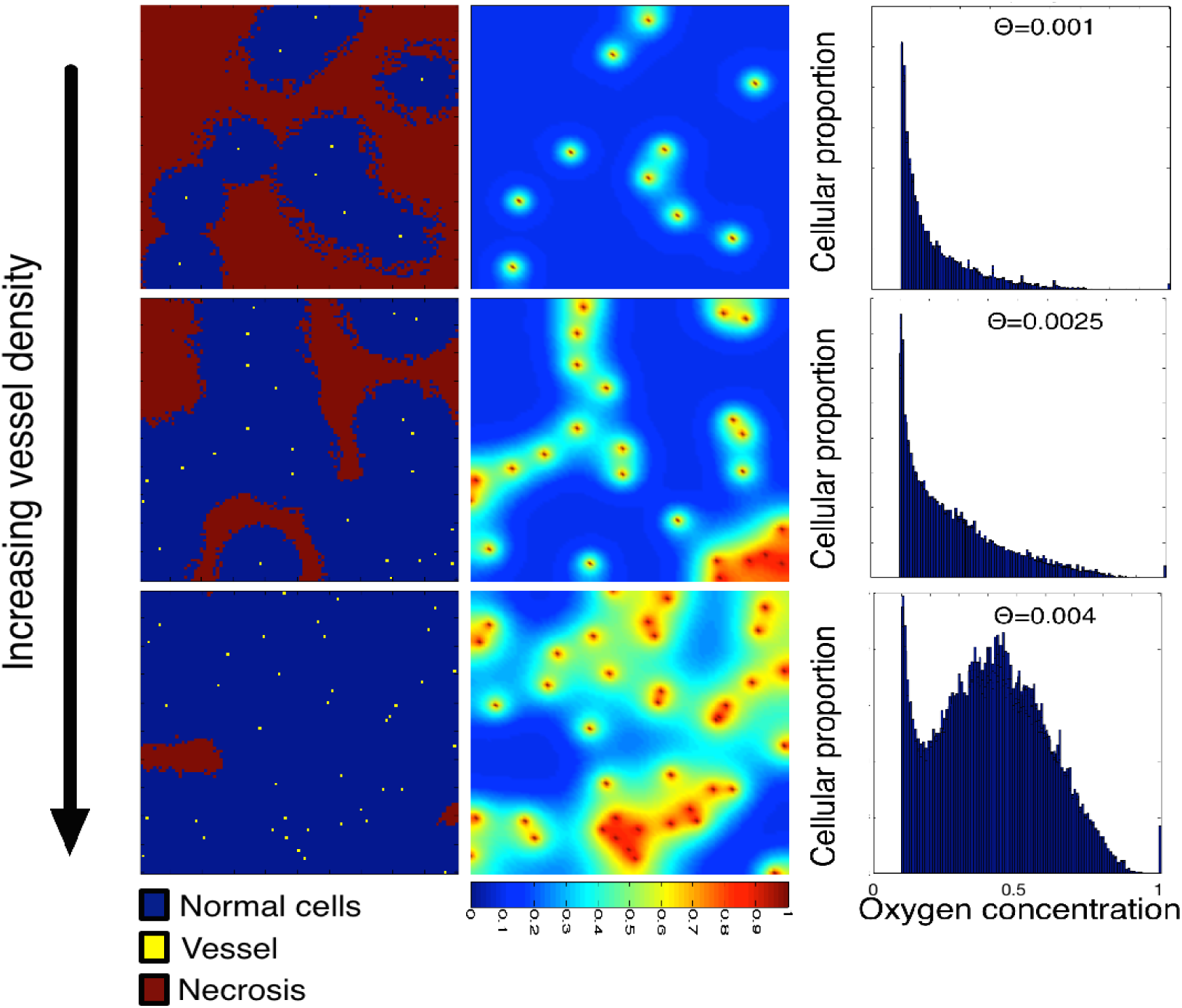
Varying vascular density affects the carrying capacity of normal tissue and the cell-oxygen concentration distribution. We show the results of normal tissue growth and maintenance as we increase the number of randomly seeded vessels from 10 (top) to 25 (middle) to 40 (bottom) on a fixed domain (100 × 100). Cells (left) and oxygen concentration (centre) are visualized. We plot the average distribution of healthy cells by oxygen concentration (right) over ten runs of the simulation with different vascular distributions but constant number of vessels. Every simulation ends in a dynamic equilibrium.

As expected, we see an increase in carrying capacity with increasing vascular density, similar to the regularly vascularized scenario, with mean cellularity (cells / domain size) increasing from 0.3445 to 0.9424, with standard deviations from 0.0208 to 0.0312. Over ten simulations in the irregularly vascularised cases, however, we find striking differences in the distribution of cell-specific oxygen concentration (Fig. 3, right). While in the regular case (Fig. 2) we find conserved cellular-oxygen distribution shapes (second and third moments), in the irregularly vascularized case we find two changes. First, even in the highest vessel density case there is a large peak at c_ap_ and second, the distribution becomes more and more skewed (to the left) as the vessel density decreases (skewness ranging from 0.2878 – 1.895).

### Vascularised tumour growth and invasion

**Tumour invasion speed changes with, and can be constrained by, vessel spacing in regular vascular architectures.** As a tumour initially invades into healthy tissue, the vasculature that it would encounter would be that of the healthy tissue. While we are ignoring the effects of cell crowding, vascular deformation and angiogenesis, it is of value to understand how the growth rate and the pattern of cancer spread would change given differing vascular spacing in an otherwise regular vasculature. To this end, we seed a series of regularly patterned vascularised domains with increasing regular vessel spacings with a single initiating cancer cell and observe the growth. Note that, to ensure initial tumour growth, we have placed an extra vessel at the centre of the computational domain with the initial cancer cell as in certain regular spacing patterns, the centre would be an area of necrosis and the simulation would end with no cancer cells.

We plot the results of these simulations in Fig. 4 along with the individual simulation growth rates. We find four qualitatively different patterns/rates of growth for different vascular densities. First, when there is high vessel density (box 1, green trace, Fig. 4), the tumour undergoes rapid growth. This approximates contact inhibited tumour growth with proliferation at the edge only. Even in this growth regime, we begin to see necrotic areas in the bulk of the tumour (box 2, blue trace, Fig. 4). The second qualitatively different regime is when the vessel density decreases to ⊝ = 0.0033, and the spacing of the vasculature is such that it has lost its ability to maintain continuous populations of cancer cells (box 3, red trace, Fig. 4). In this regime, we find that the cancer’s growth rate begins to slow, and further, that the boundary begins to become irregularly shaped as the cancer cells have to persist in regions of hypoxia long enough to place daughters in the new, adjacent, areas of increased oxygen, an event that occurs stochastically. Further reduction in vessel density to ⊝ = 0.0025 results in areas of necrosis forming in the healthy tissue and a sharp reduction in tumour growth rate (box 4, cyan trace, Fig. 4). The final scenario is defined by tissue that is so poorly vascularized that the healthy tissue cannot maintain contiguous populations, and the cancer cannot continue growing beyond the area of initial vessel oxygenation (box 5, black trace, Fig. 4), and would need to produce its own vessels to grow further.

**Figure 4.**
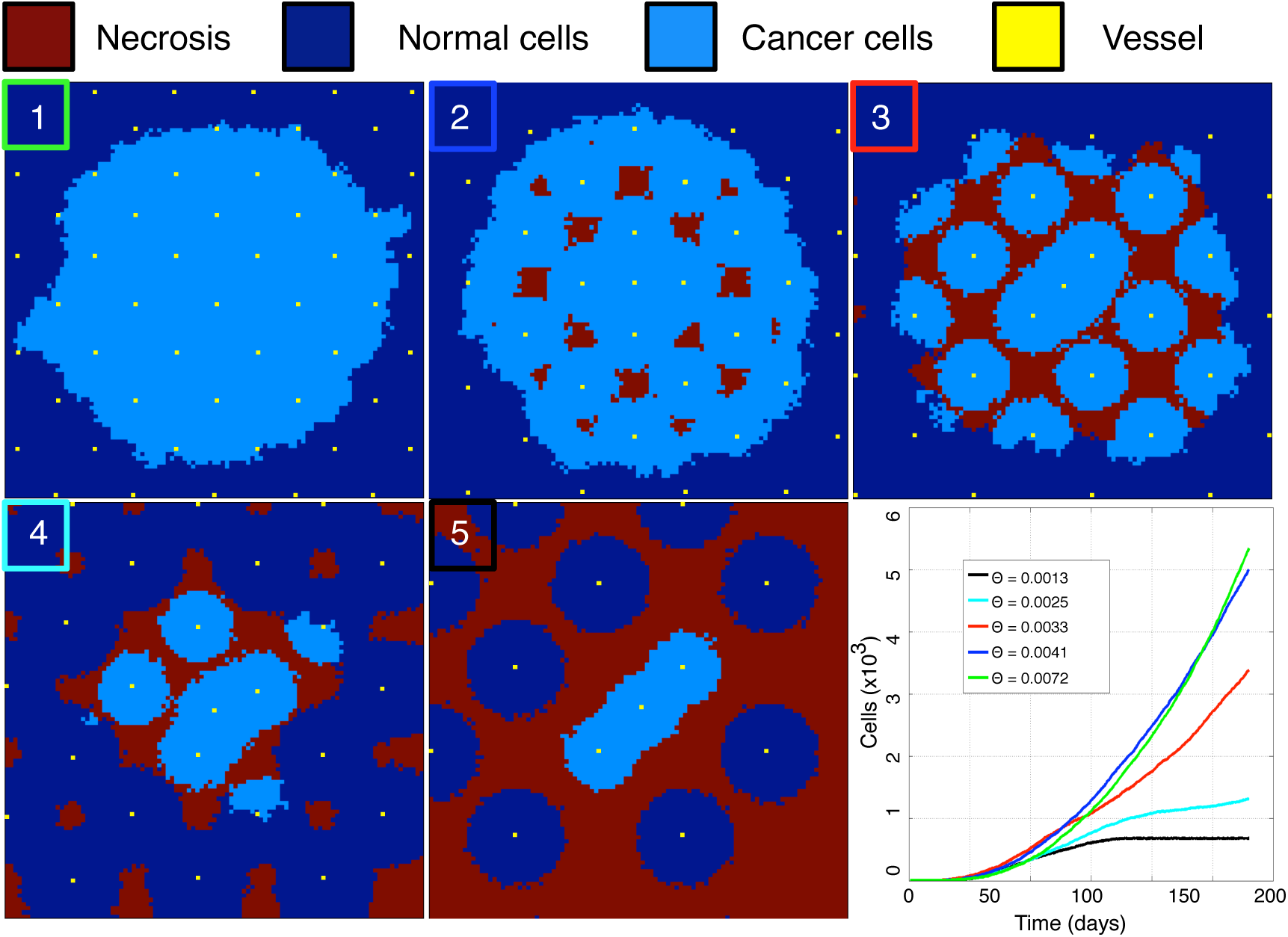
Increasing spacing between vessels slows tumour growth and creates areas of necrosis. Tumours are grown in equally sized, regularly vascularised domains with decreasing vessel density from boxes 1 to 5 (from top left: ⊝ = 0.0072, 0.0041, 0.0033, 0.0025, 0.0013). All plots show the automaton state at the final time point at time *t* ≈ 190 days. Only the smallest vessel density (0.0013 in this figure) entirely constrains growth. Growth rate over the first ≈ 190 days is summarized in the lower right.

**Tissue carrying capacity and vessel density.** We next investigate in further detail the effect of vessel density on the carrying capacity and cellular-oxygen distribution. We consider a domain of size *N* = 73 (chosen for ease of comparison for normal vessel spacing) and vary vessel number from very low until we reach saturation of cells (in this case from 3 to 236 vessels). In each simulation, we initialize the domain full of cancer cells and then allow the tissue to experience oxygen dependent birth and death until a dynamic equilibrium is reached, defined as experiencing less than a 1% change in cell number for 50 time steps. We choose this initial condition, instead of seeding the domain with a single cell, because, in this section, we are not interested in growth kinetics, but instead focus on tumour bulk equilibrium characteristics. From this point, we then record and average 100 time steps of cell and oxygen concentration data. For each vessel number, we run 500 simulations, as detailed above, with random vessel configuration and plot the result of each set of simulations and the associated standard

As expected, we see a monotonic approach to cellular saturation as vessel number increases. When we plot the standard deviation, we find that it is significantly higher in the region that is most physiologically relevant, and that these differences can reach as high as 10% (Fig. 5), making cellularity estimates based on vascular density difficult and unreliable. Further, how the different patterns represented in these families of simulations can affect the heterogeneity of oxygen within the domain, and the resultant cellular-oxygen distributions, is unknown.

**Figure 5.**
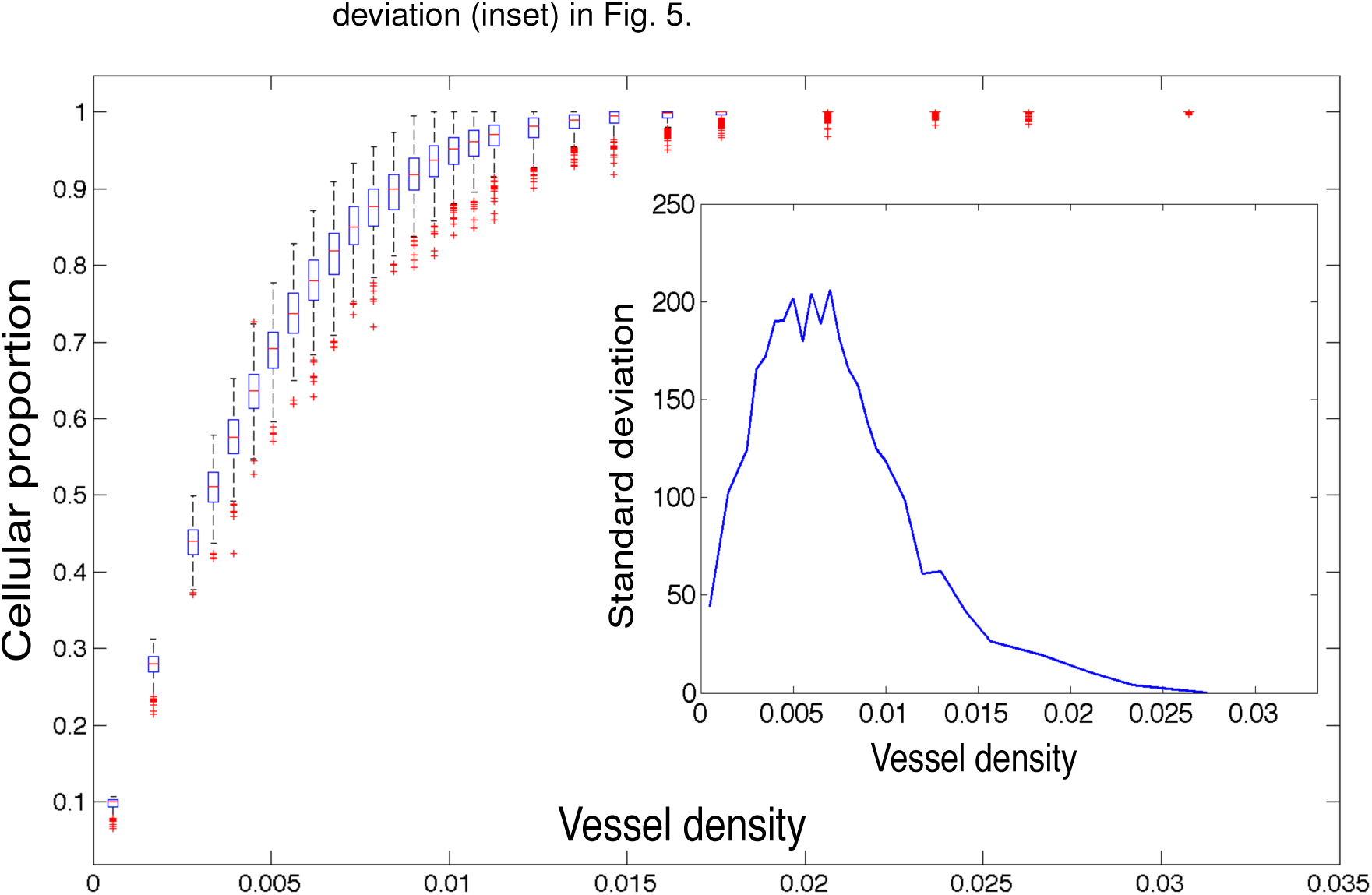
Equilibrium cancer cell density versus vessel density. We plot the results of several families of simulations seeded with equal numbers of randomly placed vessels on a fixed domain of size 73 × 73. Each data point is represented by a box which is centered on the median of 500 simulations, each of which is the average of 100 time points after dynamic equilibrium is reached. The edges of the boxes represent the 25th and 75th percentiles, the whiskers extend to the most extreme data points not considered outliers. Outliers are defined as any simulation outside approximately 2.7 standard deviations, and they are plotted as red crosses. Inset we plot the standard deviations at each vessel density.

**Effect on cellular-oxygen distribution in heterogeneous domains.** To begin to understand how the patterning of the vasculature affects the cellular-oxygen distribution within our simulations, we consider two cases from the sample of our simulations for two separate vessel densities (24 and 54 vessels). We plot and compare the vascular patterns which support the maximum and minimum cellular populations from two given vascular densities (Fig. 6). We plot the minimum cellular population represented in the left column at low (top) and high (bottom) relative vascular densities and the maximum populations in the right column.

**Figure 6.**
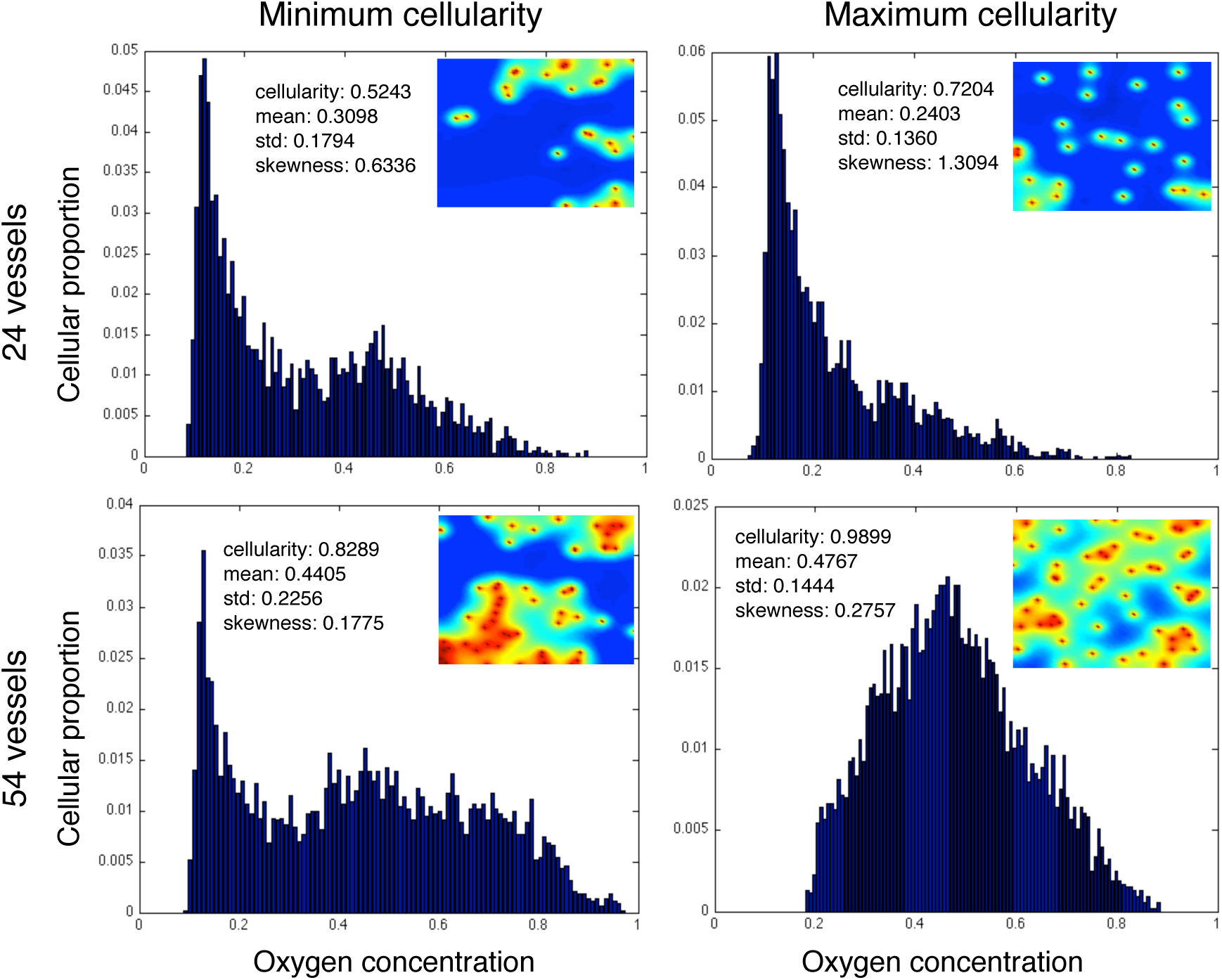
Homogeneous and heterogeneous vessel patterns with same density have very different carrying capacity and cellular oxygen distributions. We plot the equilibrium cellular-oxygen distributions and spatial oxygen distributions from the minimum and maximum cellularity examples from two representative families (24 and 54 vessels per domain) of simulations. We see nearly 20% changes in carrying capacity in favor of the more homogeneous distributions in both cases, and while the second and third moments of the distributions of oxygen distribution, the changes are highly varied from the low to high density cases.

In both minimum population cases, there is a large peak of cells near to the hypoxic minimum (*c*_ap_) as well as a smaller peak around an oxygen concentration of 0.5. In the maximum cellularity examples, however, these similarities disappear. In the low vessel density, we see that the hypoxic population is maintained, but we have lost the central, well oxygenated fraction, while in the high vessel density maximum population this is reversed, and we lose the hypoxic population and the entire population is centered around normoxia. This opposite effect (losing the hypoxic fraction in one case and increasing it in the other) would drastically change radiation efficacy, suggesting that vascular organization, not just density, could play an important role in clinical radiation therapy.

Further, we see that in each case, the maximum population is supported by more homogeneous vascular patterns, but that in the two cases the distributions are quite different, and indeed the skewness reflects this.

### The effect of heterogeneous vascular patterns on surviving cell number

As discussed in the previous section, the organisation of vessels in our model can have a significant effect on both the carrying capacity of a domain and also the resulting cellular-oxygen distribution of the cells inhabiting the domain. It is exactly these two values (cell number and surviving fraction) which influence our computation of total surviving cells, a measure critical for comparison across heterogeneous samples, and to understanding radiation efficacy at larger scales, like TCP. How these effects are governed by vessel organisation however, we have not yet elucidated. In this section we will investigate the effects of differing vascular patterns on radiation efficacy.

**Surviving cell number after radiation varies widely with vessel pattern.** To begin to understand the variability inherent in radiation effect, we calculate the number of cells surviving after a single simulated dose of 2 Gy of radiation in each simulation and plot the results as a function of vessel density (Fig. 7). We previously found that as the vessel density increases, the mean number of cells supported by the domain increases in regular domains, and this holds in irregular domains as well (Figure 10). All else being equal, this should translate into an increase in the number of surviving cells after radiation. We see that this relation holds for the low vessel density simulation families, but as the vessel density increases above 0.01, the mean number of surviving cells begins to decrease. Further, we observe, near this transition point, that the standard deviation of surviving cells within each family of simulations peaks (Fig. 7, inset). This highlights the fact that vessel density, and mean oxygen, are insufficient to predict surviving cells, therefore, in order to better predict the number of surviving cells in a domain, we require a measure of vascular organisation.

**Figure 7.**
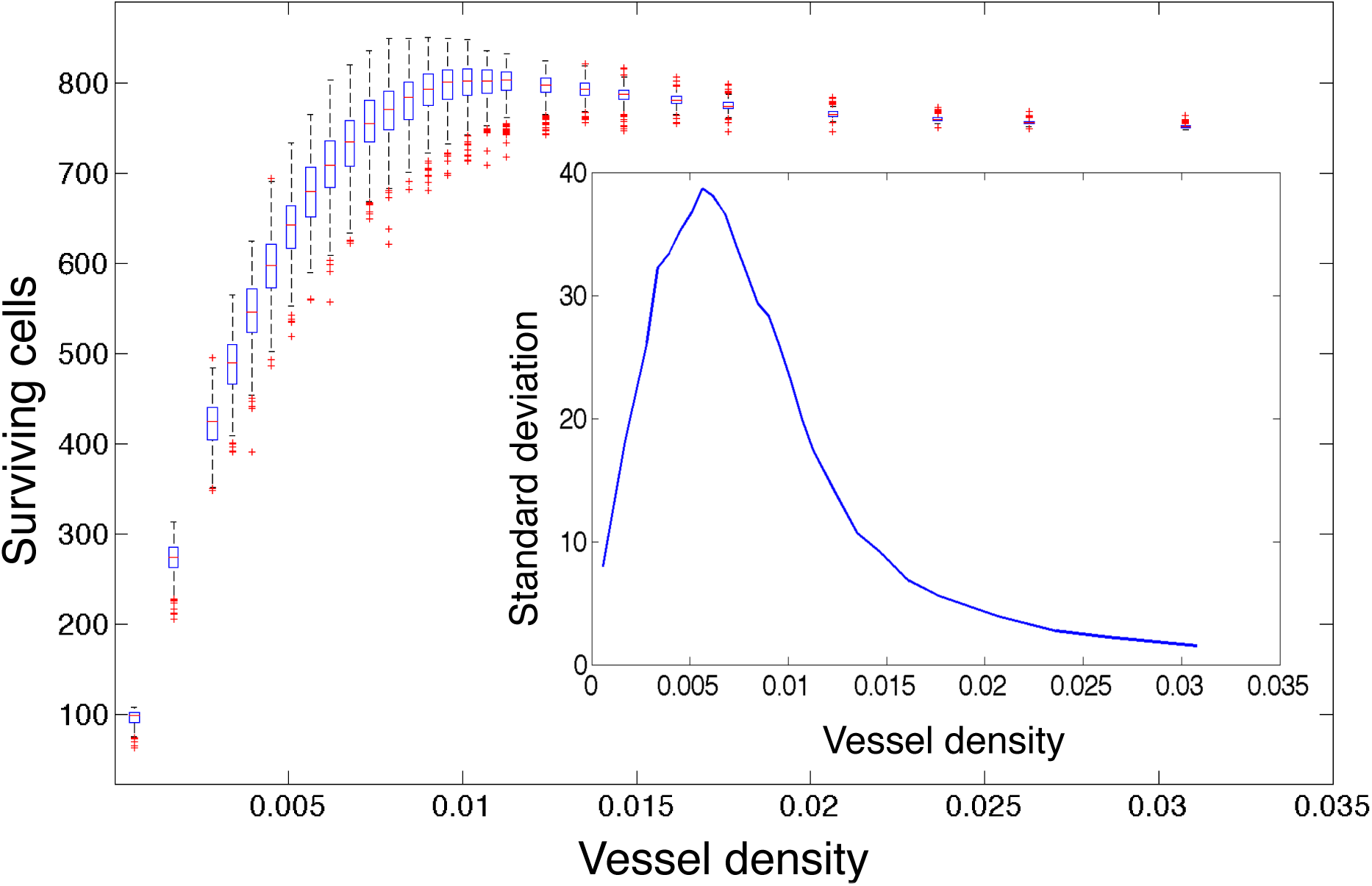
Surviving cells versus vessel density for all simulations. We plot the number of surviving cells after 2Gy of simulated radiation in each simulation as calculated using equation (5) modified by the OER from equations (6) and (7) versus the number of vessels in each case for each of the 500 simulations with constant vessel number, but random placement, on domain size 73 × 73 at dynamic equilibrium. The edges of the boxes represent the 25th and 75th percentile, the whiskers extend to the most extreme data points not considered outliers. Outliers are defined as any simulation outside approximately 2.7 standard deviations, and they are plotted as red crosses. Inset we plot the standard deviation for each family of simulations versus the vessel number.

**Vessel density is insufficient to predict radiation effect.** We have previously shown that vascular density is a strong predictor for the carrying capacity of the tissue in our model (Fig. 5). This is intuitive and, for regularly vascularized domains, as we expect in healthy tissue, is sufficient to explain not only the carrying capacity, but also other measures of the cellular distribution (e.g. the shape of the cellular-oxygen distribution). We have now shown that this measure is not sufficient when we begin to consider heterogeneously vascularized domains, as we know exist in cancer [64], where many possible spatial patterns can exhibit the same density. The mapping from cellular-oxygen distribution to surviving cells is straightforward in the simulations we have presented, as the cellular-oxygen distribution can be measured directly, and the subsequent surviving cells can be calculated using equation (13). In routinely obtained tissue biopsy specimens however, it is not possible to measuring oxygen concentrations at the cellular resolution. It would therefore be of translational value to define a metric by which cellular-oxygen distribution, or the subsequent radiation efficacy, could be inferred from a readily measurable surrogate, such as the distribution (and density) of microvessels within these samples.

### Spatial metrics of vessel organisation correlate with radiation effect

The vascular normalisation hypothesis [65] suggests that radiotherapy should be more efficacious when applied to tissues with normalised vascular beds. To test this in our model, we introduce a spatial metric by which to understand the overall level of spatial heterogeneity in our simulations.

We will utilise the variance stabilized Ripley’s *L* function [21] to measure the deviation from homogeneity in our vessel distributions. This measure, which is a function of distance, describes the average number of points within a given distance of any other point. For a complete description of this measure, see the supplemental information.

**Spatial metrics correlate directly with carrying capacity and with radiation response if vessel density is known.** In order to allow for easy correlation, we first distill this metric (which is a function of distance) into a single number by taking the mean value from 0 – 19 cell diameters (from adjacent to a distance beyond the ability to affect one another). We calculate the mean Ripley’s *L* function for all of the 500 simulations in each vessel number case and plot it against the associated carrying capacity (Supplemental Fig. 10). We find a significant negative correlation for the lower vessel densities which loses statistical significance (*p*-values not shown) as the domains begin to reach confluence for all but the minority of vessel arrangements.

Fig. 8 shows scatter plots of Ripley’s *L* function and surviving cells after radiation in 6 representative families of the 500 simulations with increasing vessel numbers. We find that at low vessel densities there is a strong negative correlation between Ripley’s *L* function and surviving cells, but this correlation changes sign at high vessel densities, explaining the counter-intuitive change in surviving fraction after ⊝ = 0.01 in Fig. 7. This suggests that in situations where the vessels are more rarefied, normalising existing vasculature could actually make radiation less effective, in contrast to the vessel normalisation hypothesis.

**Figure 8.**
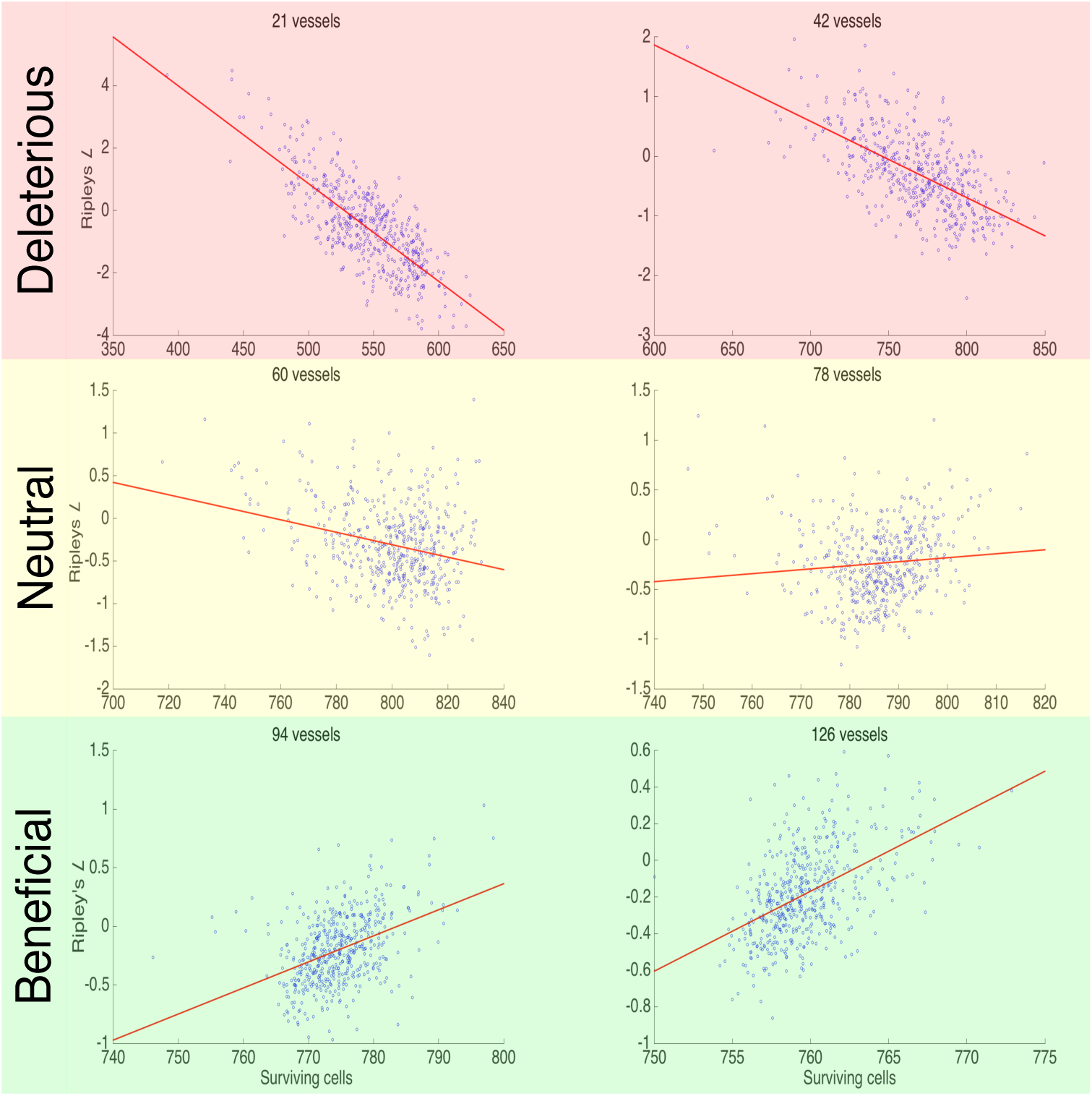
Ripley’s *L* function versus surviving cells after radiation. We plot six scatter plots showing the relationship in each of the 500 simulations represented in Fig. 5 for a given initial vessel density between cell number surviving after 2 Gy of radiation (*x*-axis) and the mean of Ripley’s *L* function (*y*-axis). We find that there is a positive correlation in the low vessel densities, and a negative correlation in the high vessel densities. All correlations are significant (p <<0.05), see Figure 11.

## Discussion

We have used an HCA model of vascular tumour growth in a planar domain to investigate the dependence of cellular oxygenation and radiotherapy efficacy on vascular density and patterning. Our results indicate that simple spatial summary statistics such as Ripley’s *L* function, which could be easily obtained from biopsy images, may predict, together with more standard measures like vessel density, radiation therapy efficacy and the effect of vascular normalisation on this. While our model is parameterised from glioblastoma data, we anticipate these results to be more widely applicable to other cancer types.

Our results corroborate those of Alarcón et al. [14], who also used a HCA approach to studying vascular tumour growth. However, our work differs in several key ways. Specifically, the vasculature in their model lies in plane rather than *en face,* a choice which was made consciously in our model to ease future translational validation with patient tissue samples. Further, we have ignored several mechanisms of competition in order to focus on defining summary measures of cellular-oxygenation status. These differences aside, Alarcón et al. found, as we did, that heterogeneity on the scale of oxygen concentration affects cell growth, both in overall speed of tumour growth and also in shape of resulting tissue. Further, they reported significant heterogeneity in steady state oxygen concentration across their domains which was dependent on vascular organisation, but this was never explored in terms of radiation effect. A similar finding was reported by Al-Shammari et al. in a biophysical model of healthy muscle tissue [17], this time in a system utilizing en face oxygen sources, as in our work. We have observed similar changes in cellular-oxygen distributions at equilibrium, and indeed, it is these changes that drive our findings concerning radiation effect.

We have seen that the assumption that mean vessel density, and subsequently mean oxygenation, can be used as a surrogate for the number of surviving cells after radiation is insufficient. Further, we found that the relationship between each of our vessel pattern measures and the surviving cells after radiation exhibits a sign change in the mid vessel density range. This change of sign in correlation means that the patterns, within a given vessel density, that take the extreme values of each measure of vascular organization can represent opposite extremes of radiation response. This suggests that if one were to perturb the vasculature, for example toward a more homogeneous distribution, one could induce opposite effects on radiation response, depending on the vessel density.

With Folkman’s discovery, in 1971, of a master tumour angiogenesis regulating factor [18], the world thought that the suggested method of blocking this factor, by which it was promised that we could ‘starve’ tumours of their oxygen supply, would dominate cancer research. It was thought that a cure for cancer would occur in a short time period. However, early trials of single agent anti-angiogenic drugs failed to produce results [20]. Later trials, with combinations of chemotherapy and anti-angiogenic drugs, however, showed promise, but even this was discordant with the leading hypothesis describing the mechanism of anti-angiogenic therapy, which was thought to entirely starve tumours of blood supply.

It was not until 2001, when Jain suggested the ‘vascular normalisation hypothesis’ [19], that these results could be understood under a single rubric. Jain suggested that anti-angiogenic drugs, instead of entirely blocking new vessel formation, worked to normalise vasculature, pruning inefficient vessels and creating a more regular lattice, thereby improving drug and oxygen delivery. More recent iterations of this hypothesis also include improvement of the efficiency of existing vessels (reviewed by Jain [65, 66]).

While this advance in our thinking has provided a way to explain the counter-intuitive results of many trials, it still does not explain why these combinations of anti-angiogenic drugs with chemotherapeutics (or radiation therapy) do not help all patients. Using our simple model system, we have observed a changing correlation with spatial measures and radiation response. This suggests that in certain cases, ‘improving’ the homogeneity of vascularisation would hurt the radiation effect, whilst in other cases it would help: a heterogeneous response to ‘normalisation’ of vascular patterning. We show that for certain cases, vessel density held constant, normal vascular patterning can respond either better or worse to radiation therapy (Fig. 8).

To translate these conclusions to the clinic would first require biological validation of these hypotheses, as well as overcoming a number of model limitations. Specifically, the assumption that all vessels are en face and all vessels are the same in terms of size and efficiency. Validation could be achieved either through *in vivo* window chamber experiments, or indirectly through examination of post-radiation surgical specimens. Model development, to include the addition of angiogenesis, biologically derived vessel geometries and vessel heterogeneity, could be used to extend these predictions to more realistic situations.

If these limitations were addressed, and validation was achieved, the technology exists currently to take advantage of this new idea. Specifically, macroscopic oxygen concentrations could be inferred from DCE MRI (or other advanced imaging) to create optimised dose plans, as suggested by Malinen et al. [7]. This information about putative vessel density could then be coupled with histologic measurements using automated localisation of vessels [26] and an algorithm to calculate Ripley’s L. These two pieces of information could then be incorporated by creating a temporally and a spatially optimised radiation plan whereby the appropriate radiation dose would be delivered before VNT to the area that would suffer after normalisation, and then the final radiation could be delivered after VNT to the area that would benefit from it.

## Acknowledgments

J.G.S. would like to thank the NIH Loan Repayment Program for support. A.G.F. is funded by the EPSRC (http://www.2020science.net) through grant EP/I017909/1. A.R.A.A. and J.G.S. gratefully acknowledge funding from the NCI Integrative Cancer Biology Program (ICBP) grant U54 CA113007 and they and P.K.M. also thank the NCI Physical Sciences in Oncology Centers U54 CA143970 grant.

## Supporting Information

### S1 Text

In the following supplementary material, we check our parameter choices using dimensional analysis and the some simple physical arguments, describe in greater detail our choice of oxygen update time step and stability requirements, perform a sensitivity analysis of our model parameters, provide more detail on the spatial statistics used in the paper and its correlation to carrying capacity and show the correlation and p-values for the plots in Figure 8.

#### Parameter estimation

We assume that normal (brain) tissue has an approximate oxygen concentration of 35mmHg [54]. This value also agrees well with an estimate of background tissue oxygen concentration of *c*_0_ = 1.7 × 10^−8^ mol cm^−2^ (4.25 × 10^−13^ mol cell^−1^) taken from Anderson and colleagues [47] after using the ideal gas law, assuming body temperature of 310K, oxygen tension of 5300Pa [54] and cell volume of 125,000*μ*m^3^.

As each of these parameters has been estimated from different sources, we will fine tune the basal oxygen consumption and physiologic vascular density for our specific case using a well-studied tumour spheroid example. We utilize the observation that the diffusion distance of oxygen to support cancer cells is approximately 10 cell diameters [67] and the information from the literature concerning the ratio of cancer to normal oxygen consumption. To estimate the baseline oxygen consumption rate then, we begin with the value *r_c_* = 2.3 × 10^−16^ mol cell^−1^ s^−1^ taken from an *in vitro* study of tumour spheroid growth [51] and then perform a virtual tumour spheroid assay (Fig. 9) to fine tune the value for our model system.

**Figure 9.**
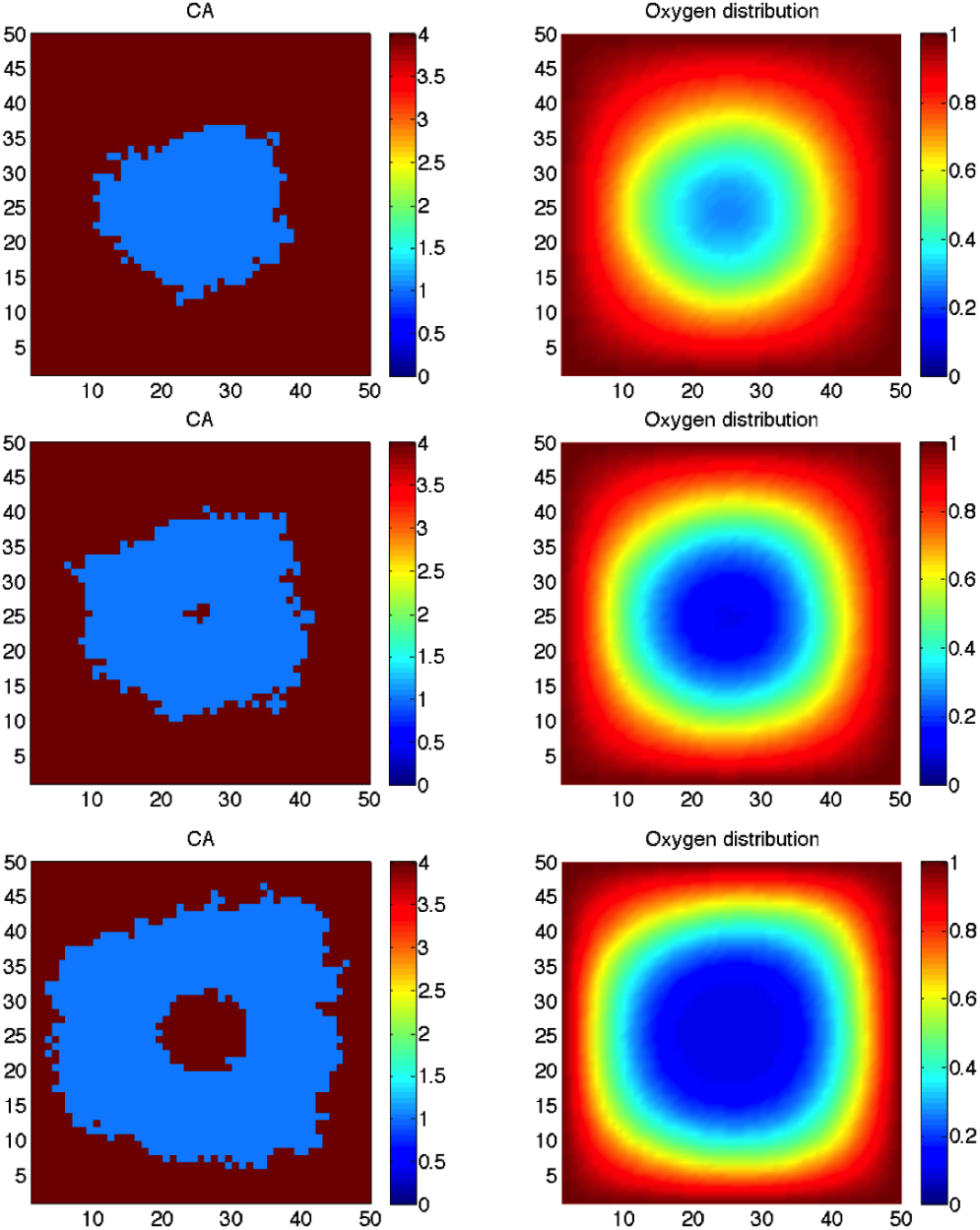
Tumour growth in an avascular domain with oxygen diffusion from the outside displays characteristics of tumour ‘spheroid’ growth. Here, an otherwise empty domain is initiated with a single cancer cell. The oxygen at the edge of the domain is set to *c* = 1. Cells (left) and oxygen concentration (right) are plotted at three time points: before the onset of central necrosis (top), initiation of central necrosis (middle) and later in progression (bottom) when a nearly constant sized proliferative ‘rim’ is observed. From this calibration, we find that the maximal oxygen uptake rate most appropriate for our model is *r_c_* = 7.5 × 10^−4^ s^−1^ which correlated with an approximate 10 cell diameter thickness.

**Figure 10.**
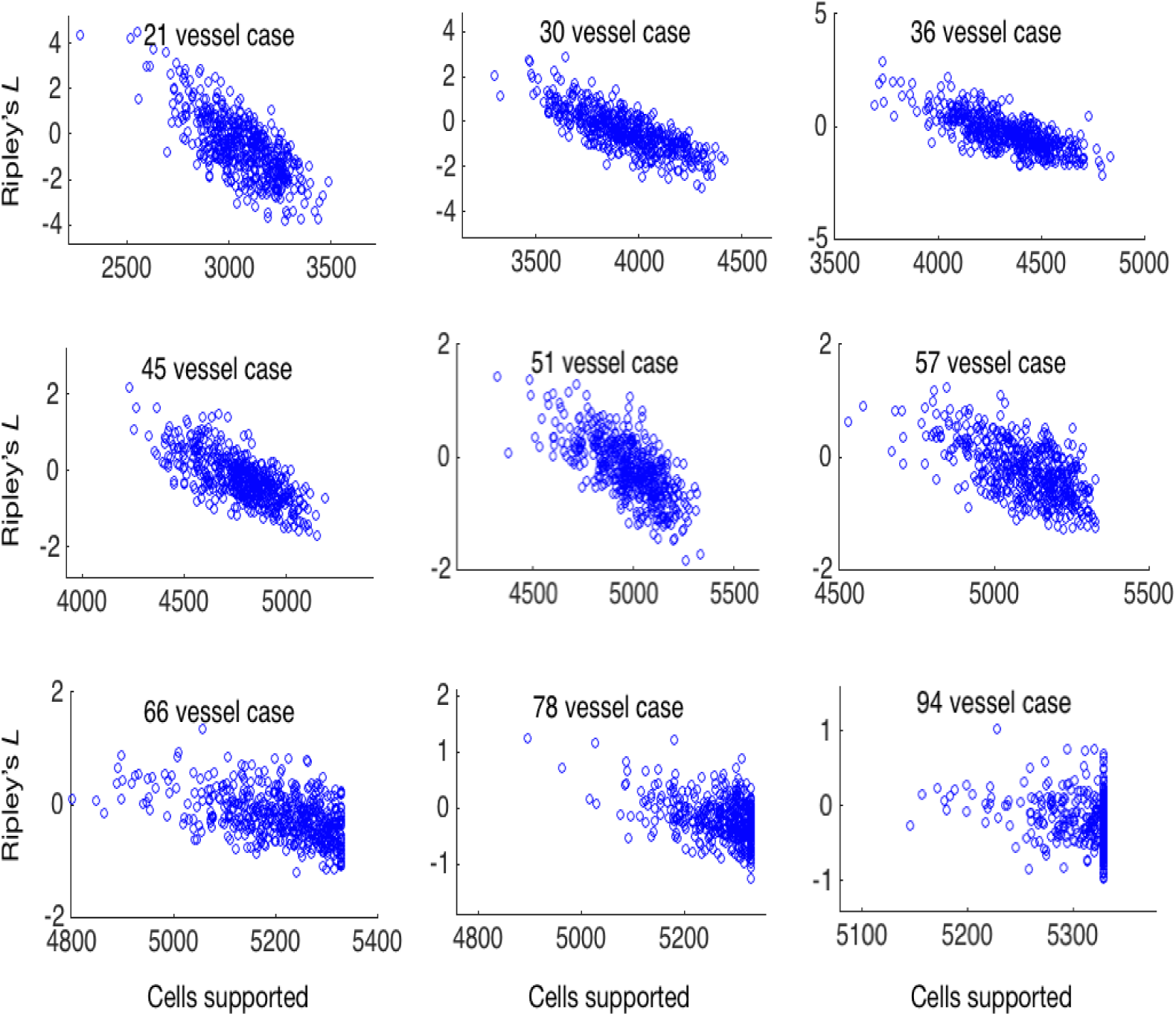
Ripley’s *L* function versus carrying capacity. We plot nine scatter plots showing the relationship in each of the 500 simulations represented in Fig. 5 for a given initial vessel density between cell number at equilibrium (*x*-axis) and Ripley’s *L* (*y*-axis). We find that there is a significant negative correlation in the low vessel densities which loses predictive capability as the domain becomes entirely filled and all of the data points align at the full carrying capacity.

**Figure 11.**
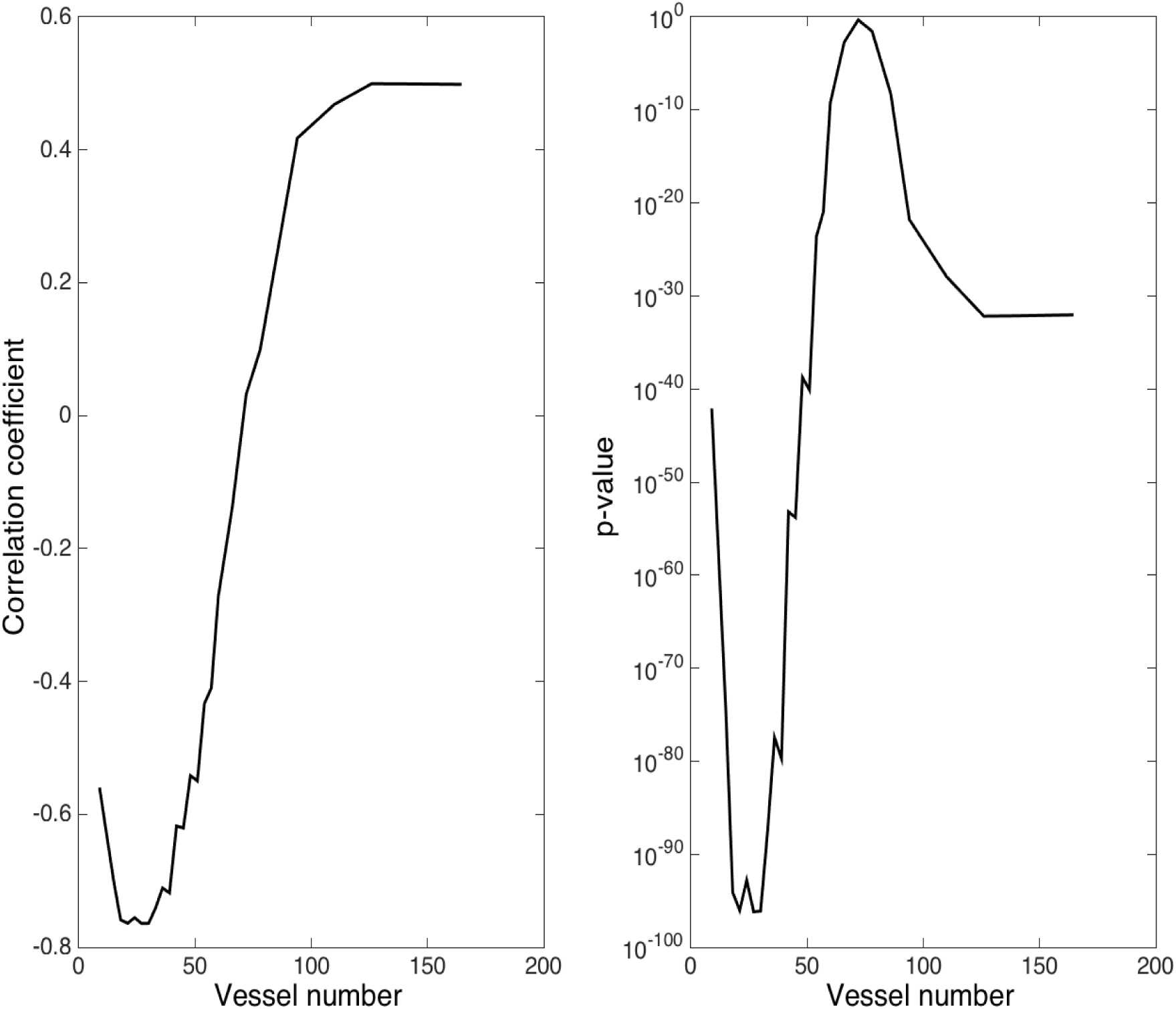
Correlation and p-value for Ripley’s *L* function vs. surviving cells after radiation. Here we plot the correlation coefficient (Left) vs. vessel density for all families of simulations and the corresponding p-value (Right). We notice that the correlation coefficient changes sign at 70 vessels, and the p-value briefly rises to insignificant (≈ 0.5) at the time of the sign change.

#### Time scales and updates

The difference in time scales between the diffusion of oxygen and the proliferation of cells is managed by updating the continuous part of the model many times per cellular time step. This can become computationally expensive in this explicit scheme, and therefore, we seek to minimize this number. However, for stability, we require Δ*tD_c_/*Δ*x^2^ <* 0.25 [68]. We therefore choose Δ*tD_c_/*Δ^2^ = 0.1. In non-dimensional parameters, we then calculate

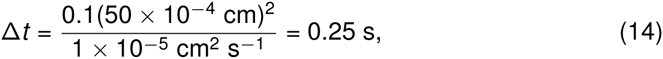
or equivalently in non-dimensional parameters

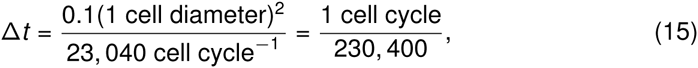
which equates to updating oxygen every 0.25 seconds, or approximately 230,400 times per cell cycle based on the parameters chosen (see Table 1). While we assume the average cell cycle time to be *τ* =16 hours, it is well known that cells in tissues are not synchronized, and also that cell fate decisions such as apoptosis are made on shorter time scales. To model this heterogeneity in cell cycle time and to more accurately match the finer time scale associated with cell death due to microenvironmental cues [69], we choose to update the cellular portion of our model 100 times per cell cycle and scale the rates for cell behaviour accordingly (reduced by a factor of 100), thereby reducing the oxygen calculations to 2,304 updates per cellular update.

**Sensitivity analysis** To assay the model for sensitivity to parameters, we measure the cellularity and cellular-oxygen distribution distributions for the regular vascularity example reported in the center panel of Figure 2, with ⊝ = 0.0027 (or a regular spacing of 14 cell diameters). From the parameter set modeled in Figure 2 (*D_c_* = 0.1, *r_c_* = 1, *K_m_* = 0.01and *c_ap_* = 0.1) we vary each parameter by ≈ three orders of magnitude and report the ranges for each mode of cellular oxygen distribution and cellularity in Table 2.

**Table 2.**
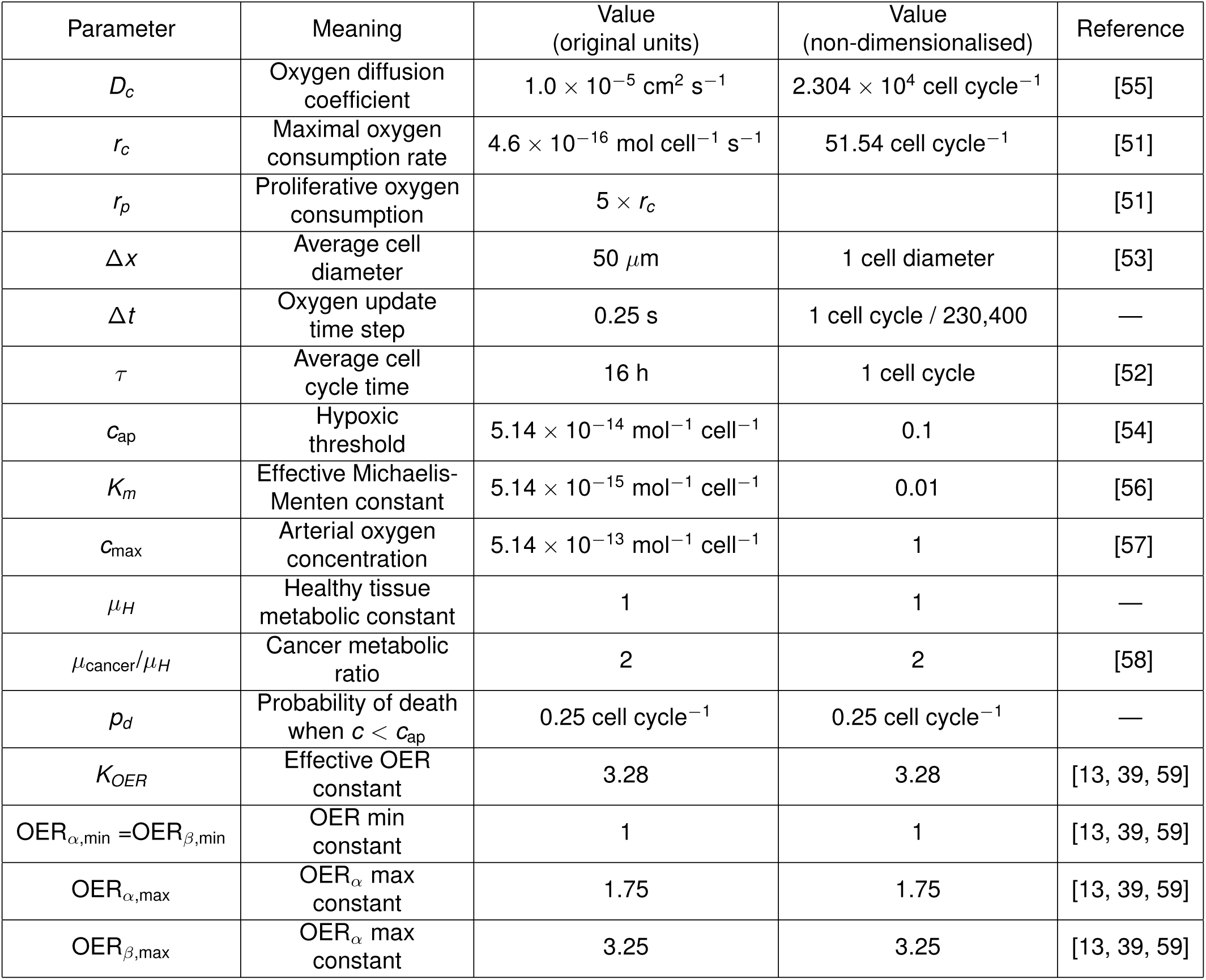
Sensitivity analysis.

As expected, increasing cellular oxygen consumption (*r_c_*) strongly influences the ability for a given vascular architecture to support cells, with greater consumption correlating with decreased cellularity. The mean oxygen experienced by cells goes down and then up slightly as the number of cells decreases drastically. Variation in the diffusion coefficient, *Dc*, strongly affects the ability for a domain to support cells, and also the mean and skewness of the resulting oxygen distribution, but affects the standard deviation relatively little. Our choice of threshold for apoptosis, *c_ap_,* intuitively has a strong effect on the cellularity, and then inversely on mean cellular oxygen as fewer and fewer cells are competing for the same oxygen. The standard deviation is affected little, and the skewness decreases at first and then increases as the number of cells becomes smaller and smaller.

#### Spatial statistics

To measure the variation away from homogeneity, we utilise a measure derived from Ripley’s *K* function. To begin, we have

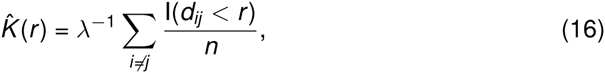
where *λ* is the average density of points in the domain, I is the indicator function which yields

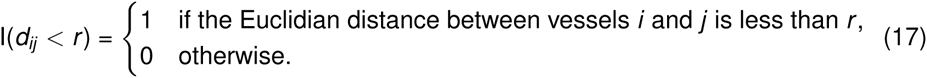

We utilize the variance stabilized version of this measure, 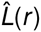 which is given by

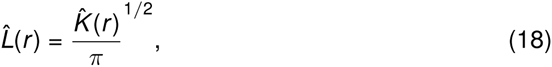
which has an expected value of 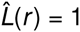 for homogeneous data. To correct for edge effects, we implement the correction suggested by Ripley [21], which changes the value of the indicator function, for points assayed within *r* of the edge, to the reciprocal of the proportion of the circle (of radius *r*) which falls outside the study area.

